# Complex chromosomal rearrangements induced by transposons in maize

**DOI:** 10.1101/2022.06.01.494422

**Authors:** Sharu Paul Sharma, Thomas Peterson

## Abstract

Eukaryotic genomes are large and complex, and gene expression can be affected by multiple regulatory elements and their positions within the dynamic chromatin architecture. Transposable Elements (TEs) are known to play important roles in genome evolution, yet questions remain as to how TEs alter genome structure and affect gene expression. Previous studies have shown that genome rearrangements can be induced by Reversed Ends Transposition (RET) involving termini of *Activator* (*Ac*) and related TEs in maize and other plants. Here, we show that complex alleles can be formed by the rapid and progressive accumulation of *Ac-*induced duplications and rearrangements. The *p1* gene enhancer in maize can induce ectopic expression of the nearby *p2* gene in pericarp tissue when placed near it via different structural rearrangements. By screening for *p2* expression, we identified and studied five cases in which multiple sequential transposition events occurred and increased the *p1* enhancer copy number. We see active *p2* expression due to multiple copies of the *p1* enhancer present near *p2* in all five cases. The *p1* enhancer effects are confirmed by the observation that loss of *p2* expression is correlated with transposition-induced excision of the *p1* enhancers. We also performed a targeted Chromosome Conformation Capture (3C) experiment to test the physical interaction between the *p1* enhancer and *p2* promoter region. Together, our results show that transposon-induced rearrangements can accumulate rapidly, and progressively increase genetic variation important for genomic evolution.

## INTRODUCTION

Enhancers are *cis*-regulatory DNA sequences that interact with their target promoters to stimulate transcription. These elements work independently of orientation and can be present near the promoter they influence, or they could act from large distances. Enhancers can be promiscuous or promoter-specific, affecting genes/promoters nearby, or passing over some genes to affect one further away (Kvon *et al*. 2014). With the discovery of *cis*-elements working from a distance, it became clear that the functionality of the genome is not only dependent on the linear DNA sequence but also on the spatial arrangement of the chromatin (Doğan and Liu 2018; Krumm. and Duan 2019). The eukaryotic nucleus has chromatin grouped into compartments of higher and lower transcriptional activity, called A and B compartments, respectively (Lieberman-Aiden, E. *et al*. 2009). Further, chromatin Topologically Associated Domains (TADs) with boundaries controlled by cohesin and CCCTC-binding factor (CTCF) play important roles in the regulation of gene expression (Dekker, J. *et al*. 2013; Nora, E.P. *et al*. 2013). CTCF is conserved within bilaterian phyla, whereas plants lack CTCF or an ortholog (Heger, P. *et al*. 2012). Hi-C maps do show compartmentalization and TADs in most plant genomes tested (Doğan and Liu 2018). Enhancers function within these domains, and structural rearrangements that change the TAD boundaries or position of an enhancer relative to their target genes can lead to dysregulation and disease (Lupianez, D.G. *et al*. 2015; Bompadre, O. and Andrey, G. 2019). With only a few well-studied examples of enhancers in plants (Weber, B. *et al*. 2016), little is known about the mechanism of their interactions with target promoters.

The maize *p1* and *p2* genes encode Myb-homologous regulators of the flavonoid biosynthetic pathway to produce red phlobaphene pigments in floral organs (Dooner *et al*. 1991; Grotewold *et al*. 1994). The *p1* gene is responsible for pigmentation in kernel pericarp, cob, and silk, while *p2* is expressed in anther and silk (Zhang *et al*. 2000; Goettel and Messing 2009). The striking red kernel phenotype specified by *p1* alleles has been used as a convenient indicator of gene expression since the earliest genetical studies in maize (Emerson 1917). The *p1* gene was one of the first loci shown to carry *Activator* (*Ac*) transposable element insertions (Barclay and Brink, 1954; Greenblatt and Brink 1962; Lechelt *et al*., 1989). The presence of 1.2 kb enhancer region near the *p1* gene was inferred from early *Ac* insertional mutagenesis studies in the allele *P1-rr4B2* (Athma *et al*. 1992; Moreno *et al*. 1992). The enhancer activity of this region was confirmed by both transient and stable transformations using the *GUS* reporter gene (Sidorenko *et al*. 1999; Sidorenko *et al*. 2000). In addition to increasing the rate of transcription, the *p1* enhancer sequence has also been shown to participate in *p1* paramutation (Sidorenko and Peterson, 2001). The *P1-rr4B2* allele has a direct duplication of 1269 bp fragment which is a part of larger 5.2 kb direct repeats which flank the *p1* gene. The 3′ 5.2 kb direct repeat contains two copies of the 1.2 kb repeat, whereas the 5′ 5.2 kb direct repeat contains one full-length 1.2 kb direct repeat, and a stable *Ds*-like element inserted into the upstream copy. The *p1* enhancer is a 405 bp fragment (Fragment 15) which lies within the 1269 bp direct duplication (Zhang, F. and Peterson, T. 2005). For simplicity, following text refers to the two copies of Fragment 15 as a single enhancer and the figures reflect the duplicate nature of the enhancer.

The maize *P1-rr11* allele has insertions of *Ac* and *fAc* (*fractured-Ac*) elements in the *p1* gene and produces a red pericarp phenotype. A derivative null allele *p1-wwB54* has white pericarp and white cob, and contains a deletion of the upstream enhancer and exons 1 and 2 of the *p1* gene (Yu *et al*. 2011). The *p1-wwB54* allele retains *p1* exon 3, the downstream enhancer, and the *Ac* and *fAc* elements in reversed orientation, separated by only 331 bp of DNA. These *Ac*/*fAc* termini can undergo frequent Reversed Ends Transposition (RET) events (Zhang and Peterson 2004; Huang and Dooner 2008; Zhang *et al*. 2009; Yu *et al*. 2011), forming a variety of alleles including deletions (Zhang, J. and Peterson, T. 2005; Zhang *et al*. 2006), duplications (Zhang *et al*. 2013), Composite Insertions (*CIs*; Zhang *et al*. 2014; Su *et al*. 2018, 2020), and inversions (Zhang and Peterson 2004; Yu *et al*. 2011; Sharma *et al*. 2021). Here we show how repeated RET events in maize can induce the formation of complex gene structures containing multiple rearrangements such as inversions, *CIs*, duplications and/or deletions. Moreover, these new structures alter the number and position of *p1* enhancer elements, thereby affecting expression of the neighboring *p2* gene.

## METHODS

### Screening and PCR

Nearly four thousand plants of genotype *p1-wwB54* heterozygous with the null allele *p1-ww[4Co63]*, were grown and pollinated with *p1-ww[4Co63]*. Ears were screened for kernels with red pericarp to obtain RET-induced rearrangement alleles. Selected red kernels were grown to obtain whole ears with red kernel pericarp. Genomic DNA was extracted from maize seedling leaves by a modified CTAB method (Saghai-Maroof *et al*. 1984) and tested for structural rearrangements using PCR (Su *et al*. 2020; Sharma *et al*. 2021). PCR was performed under standard conditions using Promega GoTaq Green Master Mix and primers specifically designed for the *p1-wwB54* sequence (Supplemental Table S1). As these rearrangements result from the activity of *Ac* and *fAc* elements, each breakpoint of any rearrangement is expected to border an *Ac* or *fAc* element. *Ac* casting (Singh *et al*. 2003; Wang and Peterson 2013) and inverse-PCR (iPCR; Ochman *et al*. 1988) techniques were used to locate these breakpoints adjacent to *Ac* and *fAc*, respectively. For visualizing PCR products, a high-efficiency agarose gel electrophoresis method was used (Sharma and Peterson 2021). PCR amplicons were sequenced by the Iowa State University DNA Sequencing Facility.

### Southern Blotting

To confirm the internal structure of the complex rearrangements, digests using *Bgl*II, *Kpn*I, *Hpa*I, and *Eco*RI restriction enzymes and their combinations were performed (data not shown for *Eco*RV and some combinations). For double digests, digestions were performed in two steps, one enzyme at a time. Whole-genome DNA was digested and loaded on an agarose gel (0.7-0.8%) run for 24-26 hrs under 35-40 volts for adequate separation of fragments. The DNA was transferred to a membrane for 24 hours, followed by probing the membranes with Fragment-15 located within the *p1* gene enhancer (Zhang, F. and Peterson, T. 2005).

### RT-PCR

Pericarps were peeled from kernels 20 days after pollination (DAP) and flash-frozen in liquid nitrogen. Two biological replicates (pericarps from 2 siblings) were pooled to extract RNA. RNA was isolated using Purelink Plant RNA Reagent, treated with New England Biolabs DNase I to remove gDNA, and then reverse transcribed to cDNA using Invitrogen™ SuperScript™ II Reverse Transcriptase kit. Protocols recommended by the product suppliers were used. Two technical replicates were used for each sample in reverse transcription reaction. Finally, PCR was performed on the cDNA using primers specific to exons 1 and 3 of *p2* to amplify the *p2* gene transcript (Supplemental Table S2). Primers specific to the GAPDH gene were used as an internal control (Supplemental Table S2).

### Chromosome Conformation Capture

A plant-specific 3C protocol was used with modifications (Louwers *et al*. 2009). Pericarps were peeled from developing kernels at 15 DAP, on ice, and added directly to a 2% formaldehyde solution made with a nuclei isolation buffer and fixed at room temperature for 1 hr in a vacuum chamber under 11.7 psi pressure. For digestion, the sample was equally divided into two tubes, 150 units of *Bgl*II were added to each, and kept overnight at 37°C while on rotation. Additional 50 units of *Bgl*II were added to each tube the next morning and incubated for 2 hrs. Slow rotations of 60 rpm were used on all steps requiring rotation. For DNA precipitation, samples were stored at -20°C. *S-adenosyl-methionine decarboxylase* (*Sam*) gene was used as an internal control in qPCR. As a control for primer efficiency differences in PCR amplification, the target regions were amplified and digested with *Bgl*II and re-ligated (Tolhuis *et al*. 2002). The DNA concentration was determined and mixed in equimolar amounts to make the control template containing all possible ligation products of the loci of interest (*p1/p2* and *Sam*). qPCR was performed using SybrGreen master mix and supplier recommended protocols.

Maize stocks are available upon request. The authors affirm that all data necessary for confirming the conclusions of the article are present within the article, figures, and tables. Supplemental figures and tables are available at figshare.

## RESULTS

Southern blot analysis using Fragment-15 (Zhang, F. and Peterson, T. 2005) allowed us to identify alleles with multiple *p1* enhancers (Supplemental Figure S1). In these alleles, rearrangement endpoints were found using *Ac* casting (Singh *et al*. 2003; Wang and Peterson 2013) and/or inverse-PCR (iPCR; Ochman *et al*. 1988) techniques. By evaluating the Southern blot results (Supplemental Figures S1, S2, and S3) together with the breakpoint sequences found with PCR techniques, the structures of different rearrangement alleles were deduced in accordance with existing models of RET (Zhang and Peterson 2004; Huang and Dooner 2008; Zhang *et al*. 2009; Yu *et al*. 2011). The presence of 8 bp TSDs at the two endpoints of each rearrangement structure confirmed their origin from *Ac* transposition (Supplemental Table S3).

The null allele *p1-wwB54* has a white kernel pericarp phenotype, and the *p2* gene is not expressed in the pericarp (Sharma *et al*. 2021). The *p1-wwB54* allele retains the *p1* enhancer downstream of *p1* exon 3; as described above, there are two nearby copies of the Fragment 15 enhancer, indicated by the two red boxes in Figure 1A (Zhang, F. and Peterson, T. 2005). Structural rearrangements that move the *p1* gene enhancer closer to the *p2* gene can induce expression of the *p2* gene in the pericarp and produce a red kernel pericarp phenotype. Secondary rearrangements are possible in many cases due to the continued presence of *Ac/fAc* termini and their ability to interact (Figure 1A). Some of the rearrangements previously studied are deletions (Zhang *et al*. 2006), inversions (Sharma *et al*. 2021), and Composite Insertions (Su *et al*. 2020). The model for the formation of Composite Insertion (*CI*) through DNA re-replication has been described previously in detail (Zhang *et al*. 2014; Su *et al*. 2020). Briefly, a *CI* arises when a transposon pair along with flanking DNA moves during replication from an already replicated part to a non-replicated region and gets re-replicated.

**Figure 1:**
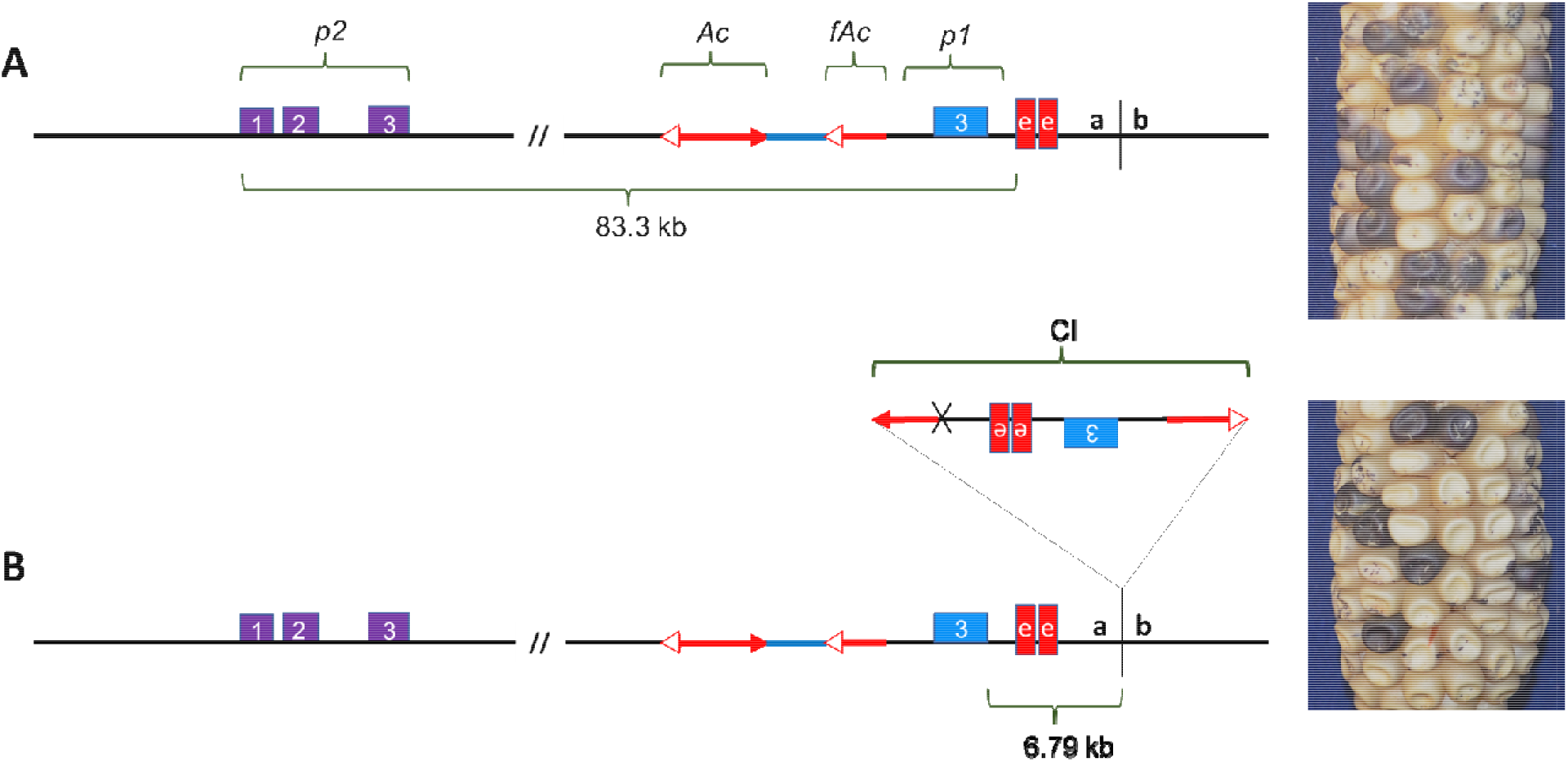
(A) *p1-wwB54:* purple boxes represent the *p2* gene with numbered exons 1, 2, and 3. The blue box is exon 3 of the *p1* gene. Red boxes indicate two copies of Fragment 15. Red arrows are *Ac* (two arrowheads) and *fAc* (single arrowhead) elements. (B) *p1-wwB54-CI* contains a *CI* at the site labeled *a/b* 6.79 kb downstream of *p1* exon 3. This *CI* contains one copy of the enhancer region and exon 3 and is also present in 5 other rearrangement alleles. Corresponding pictures on the right show colorless pericarp phenotype in both alleles. The solid and spotted purple color is due to *r1-m3::Ds* activation in aleurone caused by *Ac*-induced excision of *Ds*.

In the case of *p1-wwB54*, formation of a *CI* in the flanking downstream region created a new allele termed *p1-wwB54-CI* which contains an additional copy of the *p1* enhancer region (Figure 1B). Because the kernel pericarp phenotype of *p1-wwB54-CI* is not significantly different from the parental *p1-wwB54*, the presence of *p1-wwB54-CI* remained undetected in our materials until we characterized five independent derivative alleles. These alleles all contained a *CI* with the same internal structure inserted at the same position downstream of *p1* (Figure 1, site *a/b*). Subsequent analysis showed that our presumptive *p1-wwB54* stock contained the same *CI* at the same position, which we termed *p1-wwB54-CI*. The presence of a common 8 bp target site duplication (TSD) flanking the *CI* in all cases indicate that the *CI* insertion occurred prior to the other rearrangements unique to each case (Supplemental Table S3). The five complex rearrangements describe here appeared within five maize generations since the first isolation of *p1-wwB54* (Yu *et al*. 2011).

### Structures of complex rearrangements

Five cases with independent rearrangement structures originating from the *p1-wwB54-CI* allele were isolated. These cases contain combinations of rearrangements such as inversions, *CIs*, and deletions. In allele *p1-wwB54*-*CI* (Figure 1B), a *CI* was found to be inserted about 6.79 kb downstream of the *p1* exon 3 (Figure 1, site *a/b*). This *CI* consists of 1391 bp of *Ac* 5’ side sequence joined with 9022 bp of *fAc* and flanking region containing *p1* exon 3 and the *p1* enhancer region. These two *CI* fragments are fused at a 4 bp sequence overlap (Supplemental Table S3), suggesting they were joined by microhomology-mediated end joining (McVey and Lee 2008). In the *p1-wwB54-CI* allele (Figure 1B), two *p1* enhancers are present at 83.35 kb and 92.4 kb from *p2* 5’ end. This allele has a colorless pericarp that lacks *p2* expression like the parental *p1-wwB54* allele. The lack of expression in both alleles indicates that one or two copies of the enhancer cannot act on *p2* from this distance.

Three alleles named *SP-6, SP-7*, and *SP-12* share the same *CI* downstream of the *p1* enhancer with *p1-wwB54-CI*. In addition, these alleles have an inversion, each with a unique endpoint near *p2* (Figure 2A). The target site *x/y* in *SP-6, SP-7*, and *SP-12* is 199 bp, 257 bp, and 3041 bp away from the *p2* transcription start site (TSS), respectively, making the distances between *p2* TSS and *p1* enhancers from 4.9 kb to 16.8 kb. Target site *x/y* in each case was found to have flanking 8 bp target site duplications (Supplemental Table S3). In allele *SP-6*, the rearrangement structure leaves only 207 bp of promoter region sequence upstream of *p2*. Compared to *p1-wwB54*-*CI*, the inversions reduced the distance between *p2* and the *p1* enhancers in these cases. We expect *SP-11* (Figure 2B) to have originated from a progenitor that had the same structure as Figure 1B. The *CI* structure with *Ac* and *fAc* terminal sequences at its ends is known to be able to transpose (Su *et al*. 2020). If the *CI* excises from its *p1* location after being replicated and then inserts into a yet unreplicated region, the resulting allele will retain two copies of the *CI. SP*-*11* has the common *CI* at the *p1* downstream site, and a copy of the same *CI* is present near *p2* (Figure 2B).

**Figure 2:**
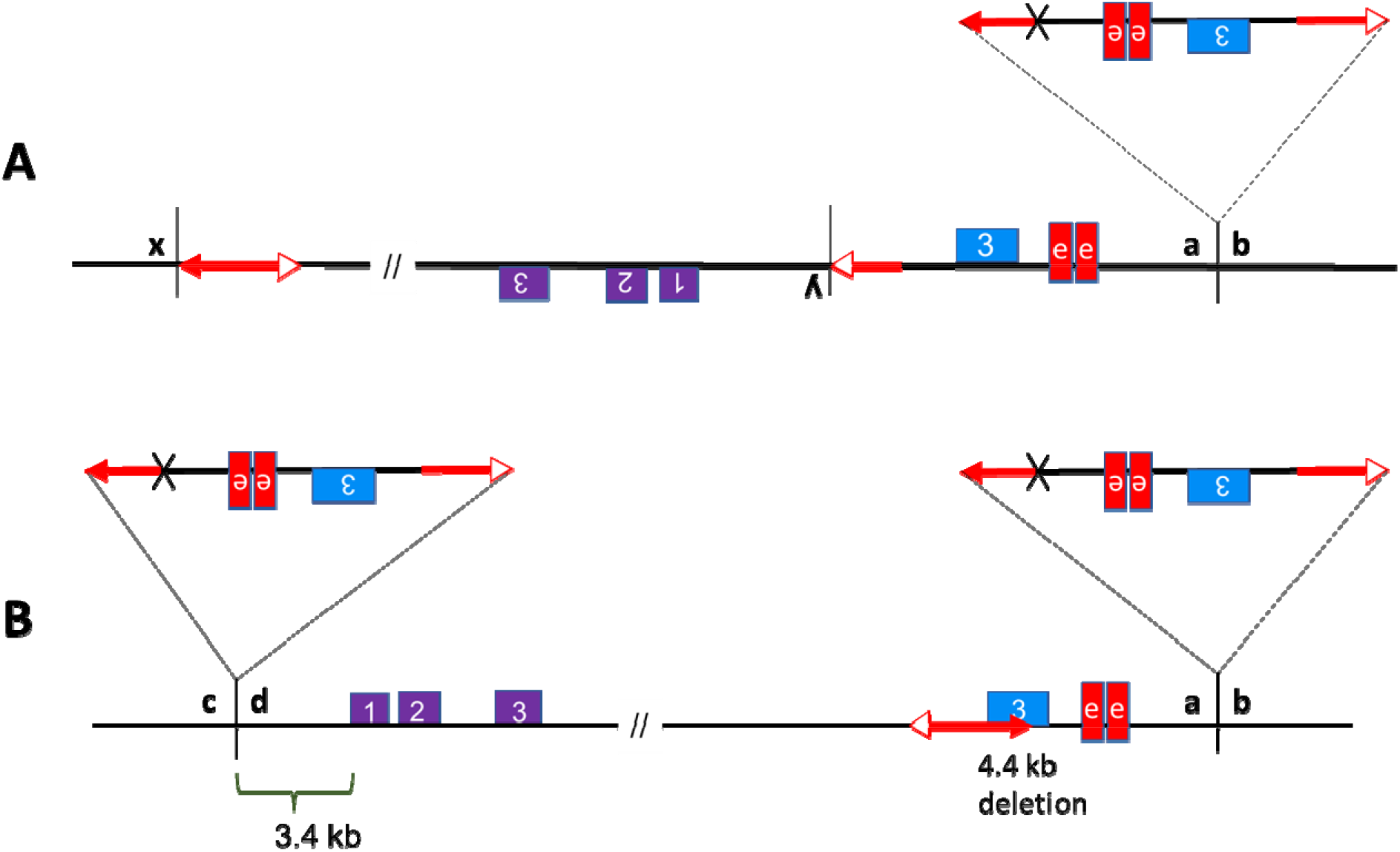
(A) *SP-6, SP-7*, and *SP-12*: These alleles have unique *x/y* inversion endpoints and share the same *a/b* insertion site for the common *CI* containing a copy of the *p1* enhancer. (B) *SP-11* has a copy of the same *CI* inserted near *p2* and an additional 4.4 kb deletion in *p1*.

In addition to the movement/duplication of the *CI, SP-11* has a deletion towards *p1* exon 3, which could result from the RET event in which the *Ac/fAc* pair inserted into *p1* exon 3, leading to deletion of the 4.4 kb fragment containing *fAc* and part of *p1* exon 3 (Figure 2B). With the two *CIs* carrying one copy of the *p1* enhancer each, *SP-11* has a total of three *p1* enhancers. The enhancer in the *CI* inserted near *p2* is at about 8.1 kb, and the two distant enhancers are at 78.95 kb and 88 kb from *p2. SP-6, SP-7*, and *SP-12* ears have a dark red pericarp phenotype, whereas the *SP-11* ear is fainter red in comparison (Figure 3). The lighter red phenotype could possibly indicate that the two distant enhancers are not involved in *p2* activation in *SP-11* same as in *p1-wwB54-CI*. RT-PCR results show that these four alleles with red pericarp have active *p2* gene expression in pericarp tissue (Supplemental Figure S4).

**Figure 3:**
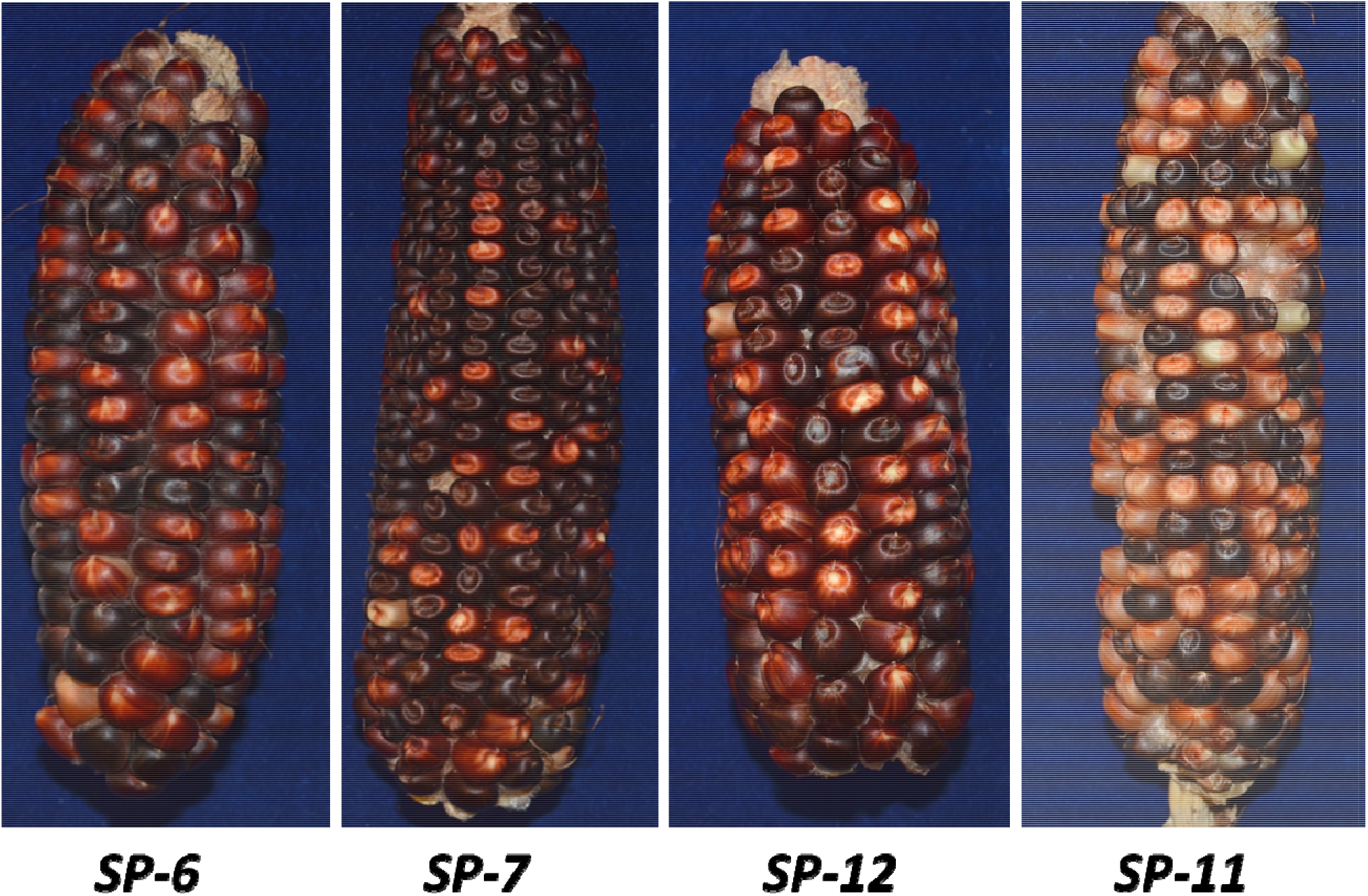
Ears of alleles with red pericarp phenotype. *SP-6, SP-7*, and *SP-12* have darker red phenotypes compared to *SP-11*.

### Progressive rearrangements in *SP-97*

The ability of multiple *Ac/fAc* elements to interact and transpose leaves many possibilities for the formation of different complicated structures. Allele *SP-97* has a complex structure with four *p1* enhancers and a dark red phenotype. Figure 4 describes a model for the origin of *SP-97*. We hypothesize that it originated after the common *CI* at the *a/b* site in *p1-wwB54* (Figure 4A, same as Figure 1B); the *Ac* and *fAc* elements present at their original position underwent reversed ends transposition into the sister chromatid at site *e/f* (Figure 4B). This resulted in a tandem direct duplication of the 17.73 kb region containing two copies of the enhancer, bringing the total to four copies (Figure 4C). The *Ac* and *fAc* elements in the recipient chromatid are also capable of generating rearrangements. A subsequent RET event with insertion of *Ac/fAc* pair at target site *x/y* 20.3 kb from *p2* caused an inversion that brought *p2* closer to the four enhancers leading to activation in the pericarp (Figure 4C, D). The size of the inverted fragment is 98.5 kb (*x* to *y* in Figure 4D). The final structure in Figure 4D lower chromatid is the *SP-97* allele containing a *CI*, a duplication, and an inversion. It would take these three RET events to form the final structure of *SP-97* leading to the red pericarp phenotype selected. The event consisting formation of the *CI* occurred prior to other two since it is present in other alleles as well. The duplication and inversion could have occurred later in a single or two different generations. The inversion would occur at last reducing the distance between the *p2* gene and the *p1* enhancers. To confirm this structure, endpoints *a/b* and *x/y* were sequenced and found to contain the flanking 8 bp target site duplications (Supplemental Table S3). The new junction created by the duplication at target sit *e/f* (Figure 4B, 4C) was also sequenced (Supplemental Table S3). The distance of the four enhancers from *p2* is about 25 kb, 34 kb, 41.5 kb, and 50.5 kb.

**Figure 4:**
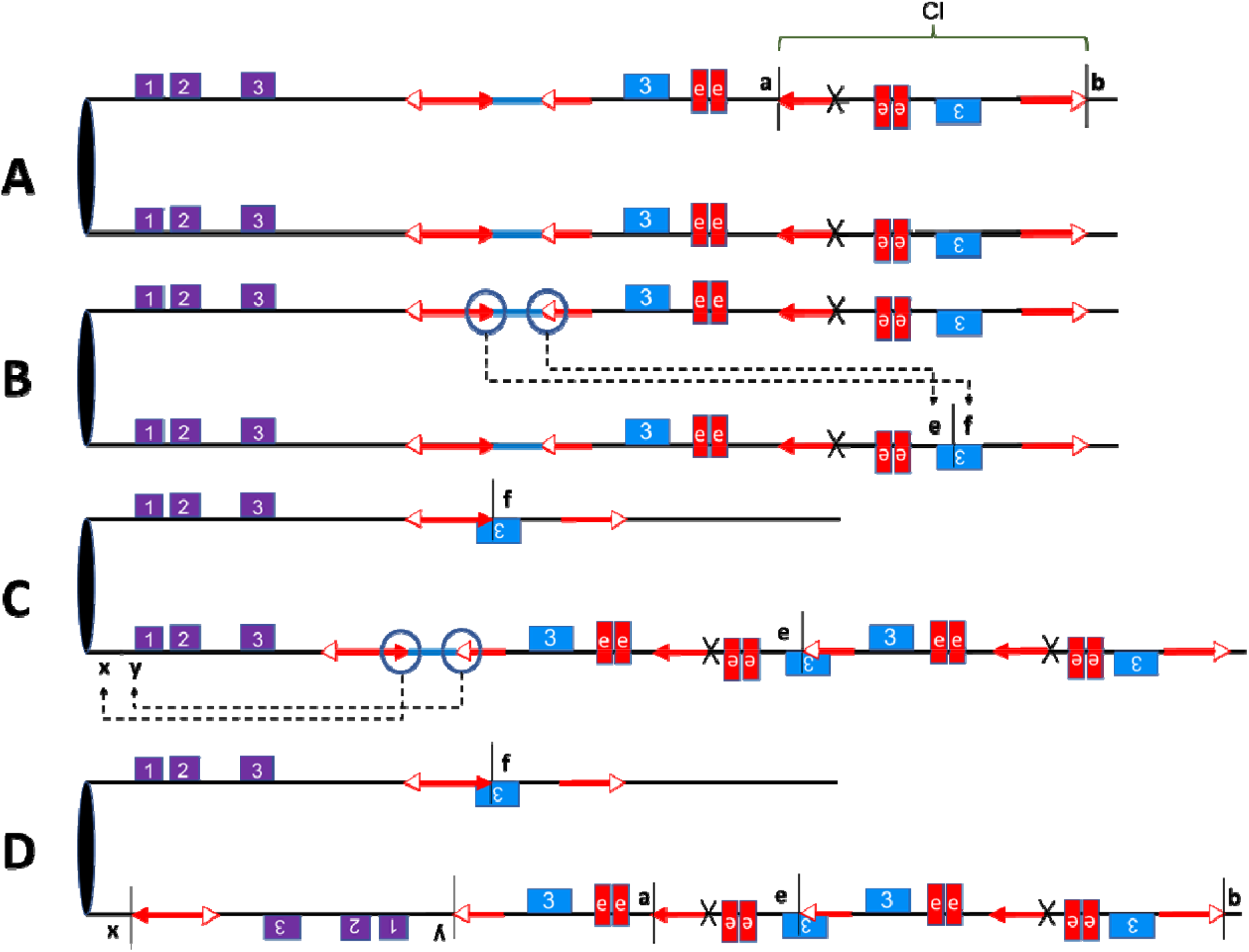
Model for the origin of allele *SP-97*: (A) Ancestral allele, which has a structure similar to *p1-wwB54-CI*. The diagram shows sister chromatids and the *CI* downstream of the *p1* enhancer at site *a/b*. (B) RET: *Ac* and *fAc* pair move from one chromatid and insert into the *e/f* target site in the sister chromatid, causing duplication and reciprocal deletion in the sister chromatids. (C) RET: The *Ac/fAc* pair on the other chromatid underwent an inversion towards *p2* inserting at target site *x/y*. (D) The lower chromatid is *SP-97*, which contains four copies of the *p1* enhancer, an inversion, a *CI*, and a duplication.

If the dark red phenotype results from multiple copies of the *p1* enhancer acting on *p2*, then the absence of some of these enhancers should affect the phenotype. To test this hypothesis, we examined ears produced by different alleles for loss of function colorless kernels. *SP-97* had some colorless/faint red kernels on the ears with dark red kernel pericarp (Figure 5A), which gave rise to the stable mutant called *SP-97M1*, which has a very faint red pericarp phenotype (Figure 5B). *SP-97* has four copies of the *p1* enhancer near *p2*. Three of the four copies are within a structure that ends with *Ac* 5’ and 3’ terminal sequences, potentially forming a 28 kb large macrotransposon, which might be capable of transposition (Huang and Dooner 2008; Su *et al*. 2020). We analyzed the structure of *SP-97M1* and found that the entire macrotransposon structure containing three copies of the *p1* enhancer has been excised; the excision site retains a modified 8-bp TSD as a macrotransposon footprint (Supplemental Table S3). The resulting *SP-97M1* allele is left with only one copy of the *p1* enhancer at 25 kb from *p2. SP-97M1* still ha some *p2* activity in the pericarp, but it is clearly lower compared to *SP-97* (Figure 6). Similarly, loss of function mutants from *SP-7* and *SP-12* (called *SP-7M1* and *SP-12M1*, respectively) were also found to have a light red pericarp phenotype (Supplemental Figure S5). In both cases, the *CI* excised out, leaving only a single copy of the *p1* enhancer near the *p2* gene. An 8 bp TSD footprint was also sequenced in *SP-7M1* (Supplemental Table S3). The lighter red phenotype in all three mutants shows that the darker red phenotype was a result of multiple copies of th enhancer present close to the *p2* gene.

**Figure 5:**
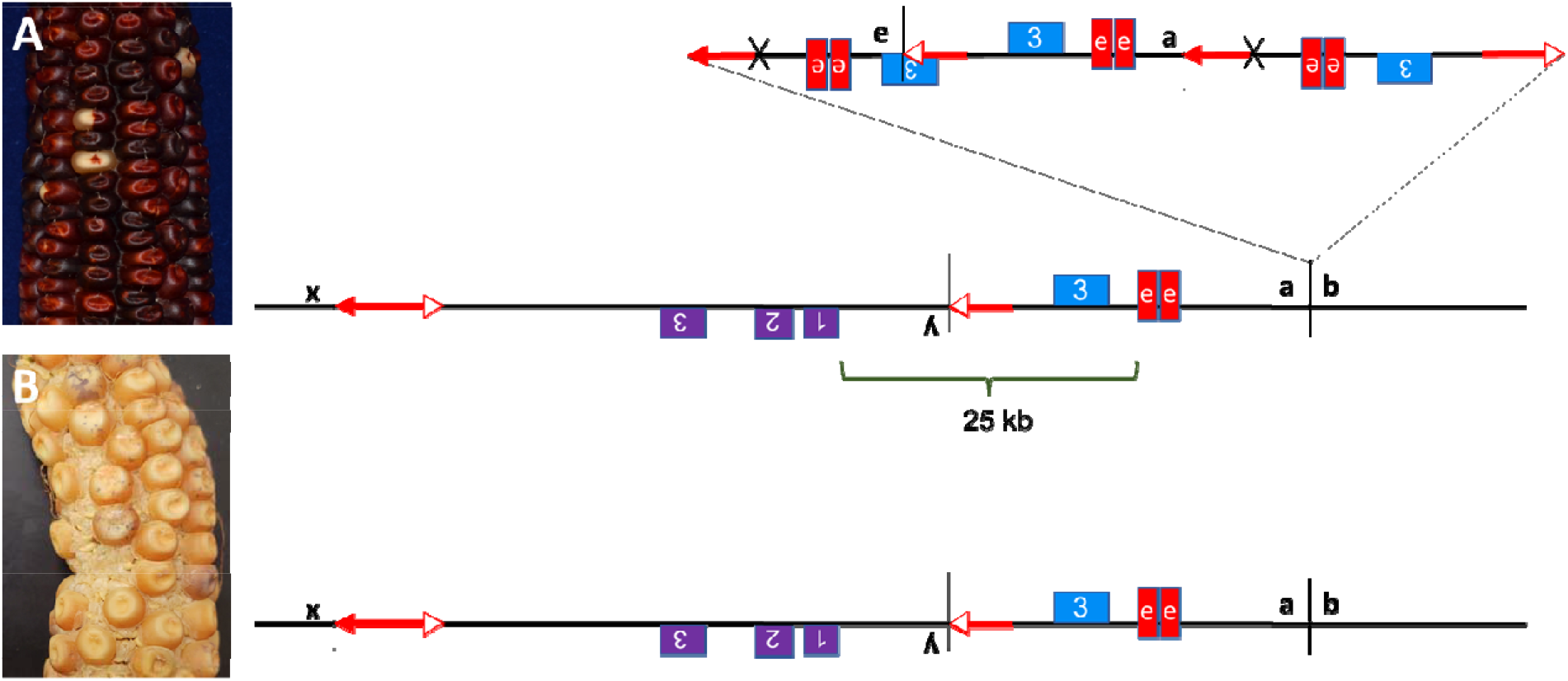
Phenotype (left) and Structure (right). (A) *SP-97* has a dark red kernel pericarp. The final structure of *SP-97* from Figure 4 is redrawn to show the 28 kb macrotransposon at site *a/b*. (B) *SP-97M1* has a light red kernel pericarp. The large macrotransposon excised out, leaving only one copy of the enhancer in the resulting allele at 25 kb from *p2*.

**Figure 6:**
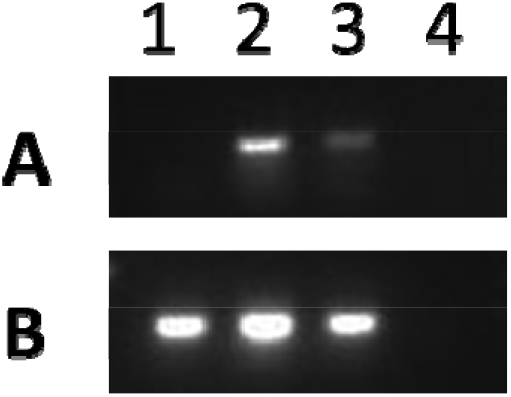
RT-PCR using RNA extracted from pericarp tissue and reverse transcribed to cDNA. Agarose gel image showing results of RT-PCR with primers from (A) *p2* exons 1 and 3, (B) GAPDH as an internal control. Lane 1, *p1-wwB54-CI*; Lane 2, *SP-97*; Lane 3, *SP-97M1*; Lane 4, negative control. *p1-wwB54-CI* lacks *p2* expression; *SP-97M1* has some *p2* expression but is lower than *SP-97*.

### Enhancer-Promoter interaction

To further test the interaction between the *p1* enhancer and the *p2* promoter, a Chromosome Conformation Capture (3C) experiment was performed on *SP-97*. First, intact nuclei were crosslinked and isolated from pericarp tissue. Then the nuclei were digested with *Bgl*II and re-ligated. After reversal of crosslinking, the resulting 3C DNA consists of new DNA molecule formed by ligation of interacting loci. These new molecules were tested using primers specific to *Bgl*II fragments around *p1/p2* region (Supplemental Table S4). Due to the presence of many repetitive DNA sequences arising from retroelements within a 100 kb region around *p2*, many *Bgl*II fragments could not be tested. The interaction frequency between the fragments containing *p1* enhancers and the region around *p2* was determined (Figure 7). In Figure 7, the fragment labelled VIII has the same sequence as fragment X, and fragment IX is the same as fragment XI. This is due to the duplication event in the origin of *SP-97* (Figure 4B, 4C). The primer specific to fragments VIII and X was used as an anchor and tested against primers specific to fragments I to VII. The peak in the figure corresponds to fragment III containing the promoter region and exons 1 and 2 of the *p2* gene, and lower interaction frequency in other nearby fragments indicates the interaction is specific between the *p1* enhancer and the *p2* gene.

**Figure 7:**
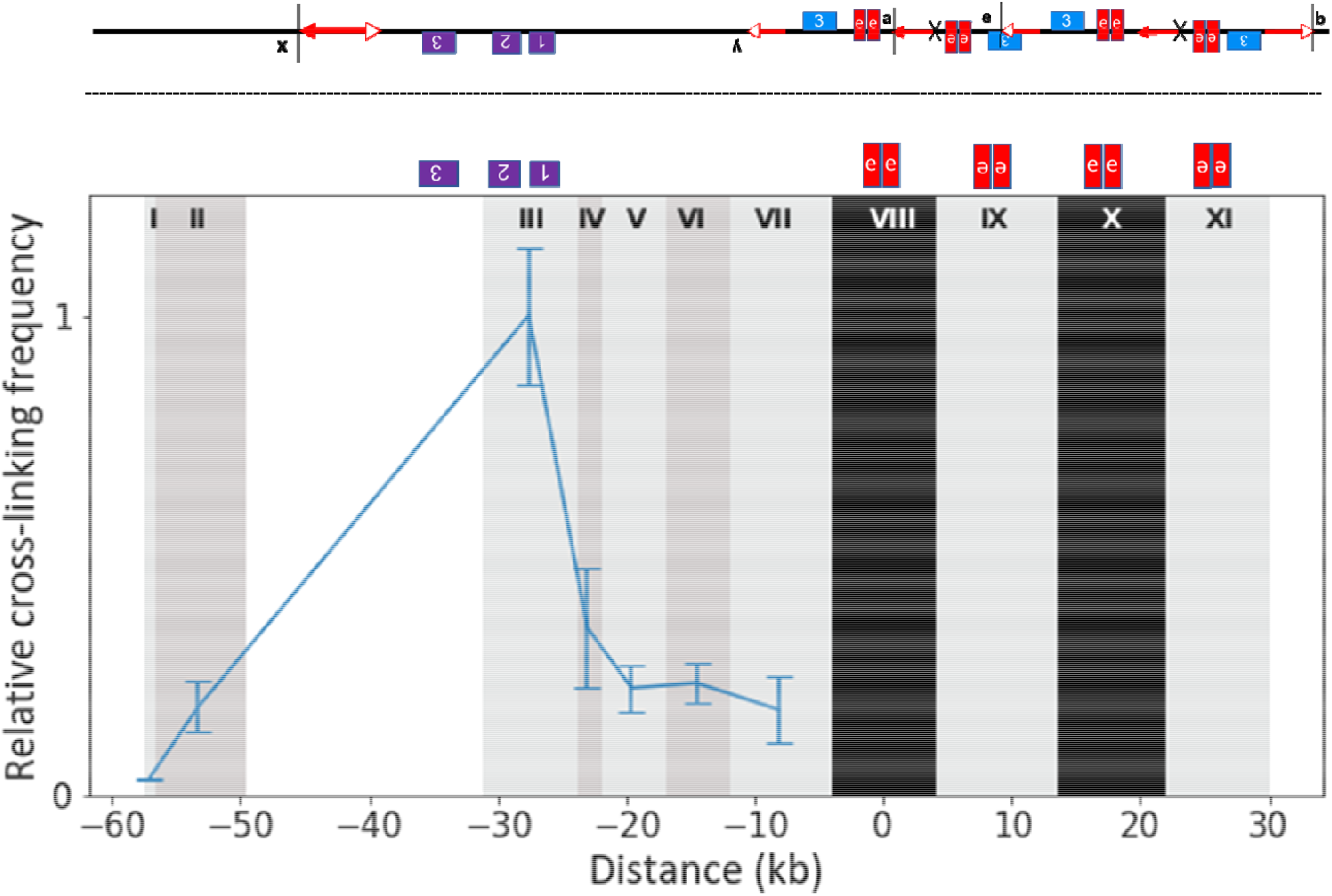
(Top) The structure of SP-97 from Figure 4D. (Bottom) Relative cross-linking frequency at *p1/p2* locus in *SP-97*. The vertical shaded columns indicate the location of *Bgl*II fragments, numbered with Roman numerals. The anchor fragments are shaded in black. The blue line shows the relative crosslinking frequency of fragments tested against the anchors. Error bars indicate the standard error of the mean of 3 samples. The y-axis is cross-linking frequency, and the x-axis is the distance in kb. Location of *p1* enhancers and *p2* gene are shown above the graph.

## DISCUSSION

### Progressive rearrangements form complex alleles

Complex Chromosomal Rearrangements (CCRs) are defined in the human genetics literature a structures consisting of more than two breakpoints and involving two or more chromosomes (F. Pellestor *et al*. 2011). CCRs are clinically important as they are involved in a number of abnormal phenotypes such as recurrent miscarriages (Daniela Giardino *et al*. 2009), mental retardation (J. R. Batanian and M. S. Eswara, 1998), and congenital malformations (Stefan Vermeulen *et al*. 2004). In addition to disease-causing variation, CCRs are also important for the formation of loci with adaptive benefits in both animals and plants. Fibrinogen locus in humans, which is a major clotting factor (Jeffrey A. Kant *et al*. 1985), and the *sh2*□*R* allele of the maize *shrunken*□*2* locus which gave rise to the sweetcorns (Vance Kramer *et al*. 2015; Ying Hu *et al*. 2021) are two examples of alleles originating from multiple rearrangements leading to a complex structure. The different *p2* alleles we present here are also examples of CCRs in plants, except for the involvement of a single chromosome in their formation. We show that a pair of DNA transposable elements can form CCRs by recurrent alternative transposition events, leading to the formation of complex alleles that can alter gene expression.

Previous studies have shown that alternative transposition of *Ac/Ds* elements can generate genomic rearrangements (Zhang *et al*. 2009; Yu *et al*. 2011). Even a single rearrangement event can have a dramatic effect on gene expression (Sharma *et al*. 2021). In this study, we used the red pericarp phenotype as an indicator of *p2* gene activation. We screened for red kernels to identify rearrangements caused by the movement of a pair of DNA elements (*Ac/fAc*). We show that these elements remain active and capable of undergoing transpositions causing progressive rearrangements. We identified CCRs with *p2* activity, but it is possible for other CCRs to occur without activation of *p2*. So, although we describe only five CCRs among 4,000 ears, the total number of CCRs is likely much higher. The five CCR alleles described here consist of multiple rearrangement events leading to the activation of the *p2* gene due to the presence of the *p1* enhancer in close proximity. In addition to the enhancer being able to interact with the target gene, the loss of function in *SP-97M1, SP-7M1*, and *SP-12M1* alleles with a decrease in the number of copies of the enhancer indicates that multiple enhancers work together to induce the comparatively darker red phenotype in *SP-97, SP-7*, and *SP-12*.

RET-induced DNA re-replication can generate *CIs* that act as *Ac*-macrotransposons (Zhang *et al*. 2014; Su, W. *et al*. 2020). *Ac*-macrotransposons have been found in several different maize lines highlighting their role in genome evolution (Wang, D. *et al*. 2022). The *SP-11* allele is an example of the reinsertion of a 10.3 kb *CI* containing the *p1* enhancer near the *p2* gene. The retention of the macrotransposon at its original location together with the insertion of a second copy shows that these macrotransposons can increase in number, as previously proposed for *Ac* element transposition (Greenblatt and Brink 1962; Chen *et al*. 1987). In addition to the 10.3 kb macrotransposon present in all the cases discussed in our results, we present the example of *SP-97*, which contains a large macrotransposon of 28 kb size. The mutant *SP-97M1* shows this macrotransposon is able to excise. *Ac/fAc* termini in reverse or direct orientation have been known to cause chromosome breaks at a frequency inversely proportional to the distance between the interacting termini (Yu *et al*. 2010). Both *SP-12* and *SP-97* were found capable of chromosome breakage, although breakage is much more frequent in *SP-97* (Supplemental Figure S6), likely due to the presence of numerous closely-spaced *Ac/fAc* termini.

### Effects of multiple enhancers

There is clearly a limited distance up to which the *p1* enhancer can influence *p2* gene expression. The two cases which have the *p1* enhancers more than 80 kb away from *p2*, the parental *p1-wwB54* allele with one *p1* enhancer and *p1-wwB54-CI* with two *p1* enhancers, both have colorless pericarp. But all alleles with active *p2* expression in the pericarp have at least one enhancer within 25 kb of *p2*, indicating that the distance at which a single enhancer can no longer interact with *p2* is somewhere between 25 kb and 80 kb. Among cases with a single *p1* enhancer, the pericarp color gets lighter with an increase in distance (Figure 8), indicating the interaction is distance-dependent. A positive correlation between gene expression and enhancer proximity has also been reported at a genomic scale (Downes, D. J. *et al*. 2021) and a single locus (Nolis, I. K. *et al*. 2009). Although enhancers are known to work from large distances, these long-range interactions are enabled by facilitating mechanisms such as chromatin looping and transcription factors that help the enhancer to reach the target promoter (Wulan Deng *et al*. 2012; Bartman *et al*. 2016). In addition to these facilitating mechanisms, an enhancer might also have an intrinsic range in which it can interact with its target promoter. In a 2009 study using transgenic HeLa cells, Nolis, I. K. *et al*. show the IFN-β enhancer activates transcription only up to a distance of 560 bp but with the addition of binding sites for Sp1 and CCAAT enhancer binding protein (C/EBP) transcription factors, the range increases to at least 2325 bp. In our case, *SP-97M1* has a lighter red phenotype with the single enhancer at 25 kb, and *SP-97* has a darker red phenotype with additional enhancers at distances of 34 kb, 41.5 kb, and 50.5 kb, which are all larger than 25 kb (Figure 4, 5), suggesting that presence of multiple enhancers can increase the maximum distance at which productive enhancer-promoter interactions can occur. If increasing the enhancer number strengthens the enhancer-promoter interaction, it is possible that it could also disrupt the 3D chromatin structure (Chakraborty, S. *et al*. 2022).

**Figure 8:**
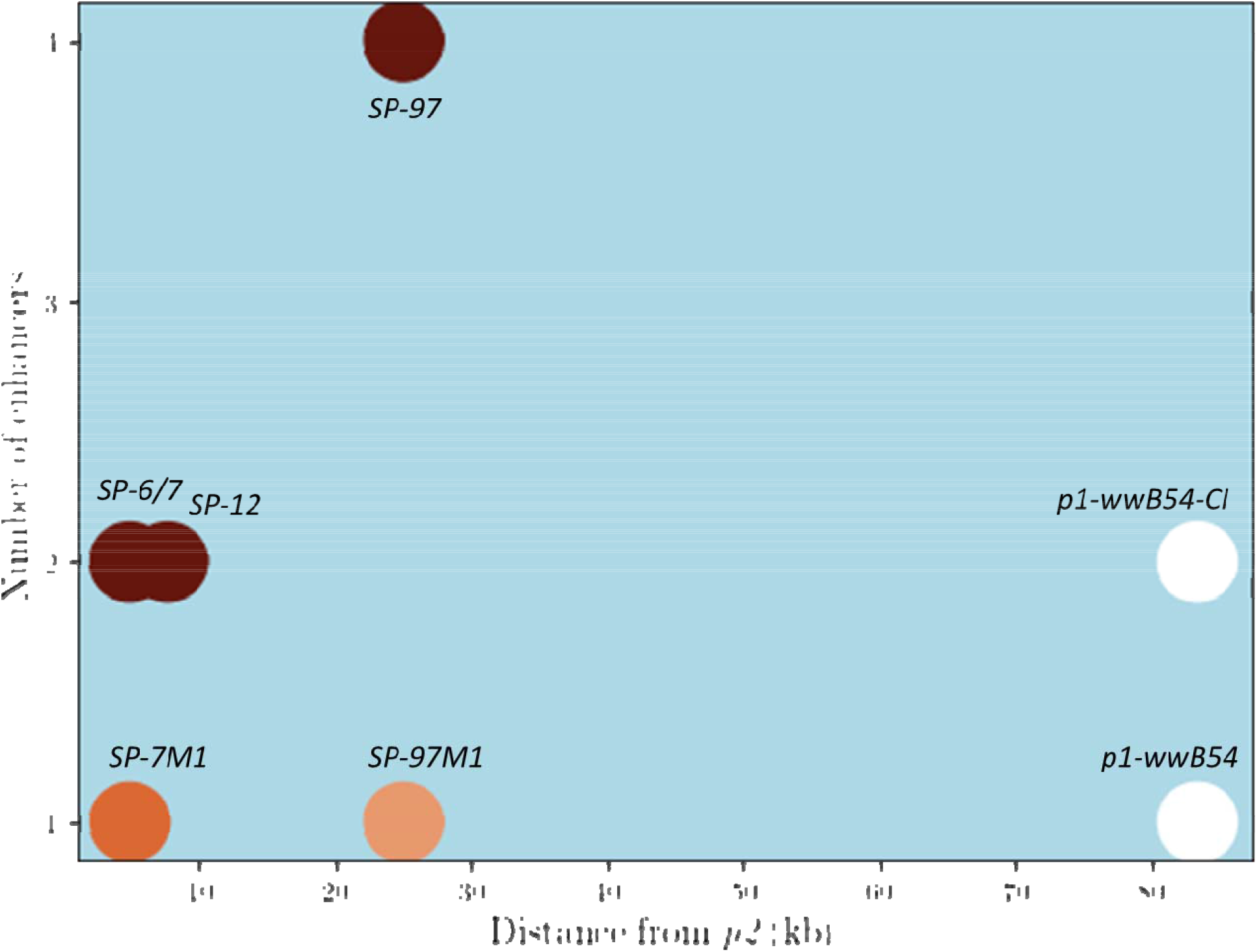
Relationship between enhancer number, distance from *p2*, and pericarp phenotype. The circles represent different alleles and their pericarp color, the x-axis is the distance of the nearest enhancer in kb from *p2*, and the y-axis is the number of enhancers. The darker red phenotype seems to have a positive correlation with the number of enhancers and a negative correlation with the distance from the target gene.

Long-range transcriptional *cis*-regulatory elements (CREs), including enhancers, are widespread in the maize genome (Ricci, W.A. *et al*. 2019). A hepta-repeat present at 100 kb upstream from the *b1* gene and distal *cis*-element (DICE) present at 140 kb upstream of the *BX1* gene both function as enhancers for their respective target genes. In both cases, alleles containing multiple copies of the enhancer have higher expression compared to their single enhancer counterpart (Louwers, M. *et al*. 2009; Zheng, L. *et al*. 2015). In our system, the pericarp phenotype is darker red in cases with multiple copies of the enhancer compared to the lighter red phenotype in cases with a single copy of the enhancer near the *p2* gene (Figure 8). *SP-97* with 4 enhancers has higher expression compared to its single enhancer mutant *SP-97M1* (Figure 6). A similar correlation between enhancer number and expression has been reported in transgenic grape and tobacco plants (Li, Z. T. *et al*. 2003). Having multiple copies of an enhancer is known to increase expression in mammalian systems as well (Downes, D. J. *et al*. 2021). Although a change in enhancer number and position can cause misexpression (Will, A. J. *et al*. 2017), in some contexts having multiple enhancers can be advantageous as it can provide robustness against disease-causing mutations (Wang, X. and Goldstein, D. B. 2020; Osterwalder, M. *et al*. 2018). In summary, we show that RET-induced rearrangements can change enhancer copy number and position, fueling *cis*-regulatory variation vital for genomic evolution.

## Data Availability

Maize genetic stocks are available upon request. Supplemental material available on figshare.

## Acknowledgements

We thank Terry Olson and Libuse Brachova for technical assistance. We thank Harry T. Horner, Mohan Gupta, Allison Birnbaum, Erica Unger-Wallace and Weijia Su for their assistance and contributions to the project.

