## Supplemental Figures and Tables for "Complex chromosomal rearrangements induced by transposons in maize"

**Supplemental Information**

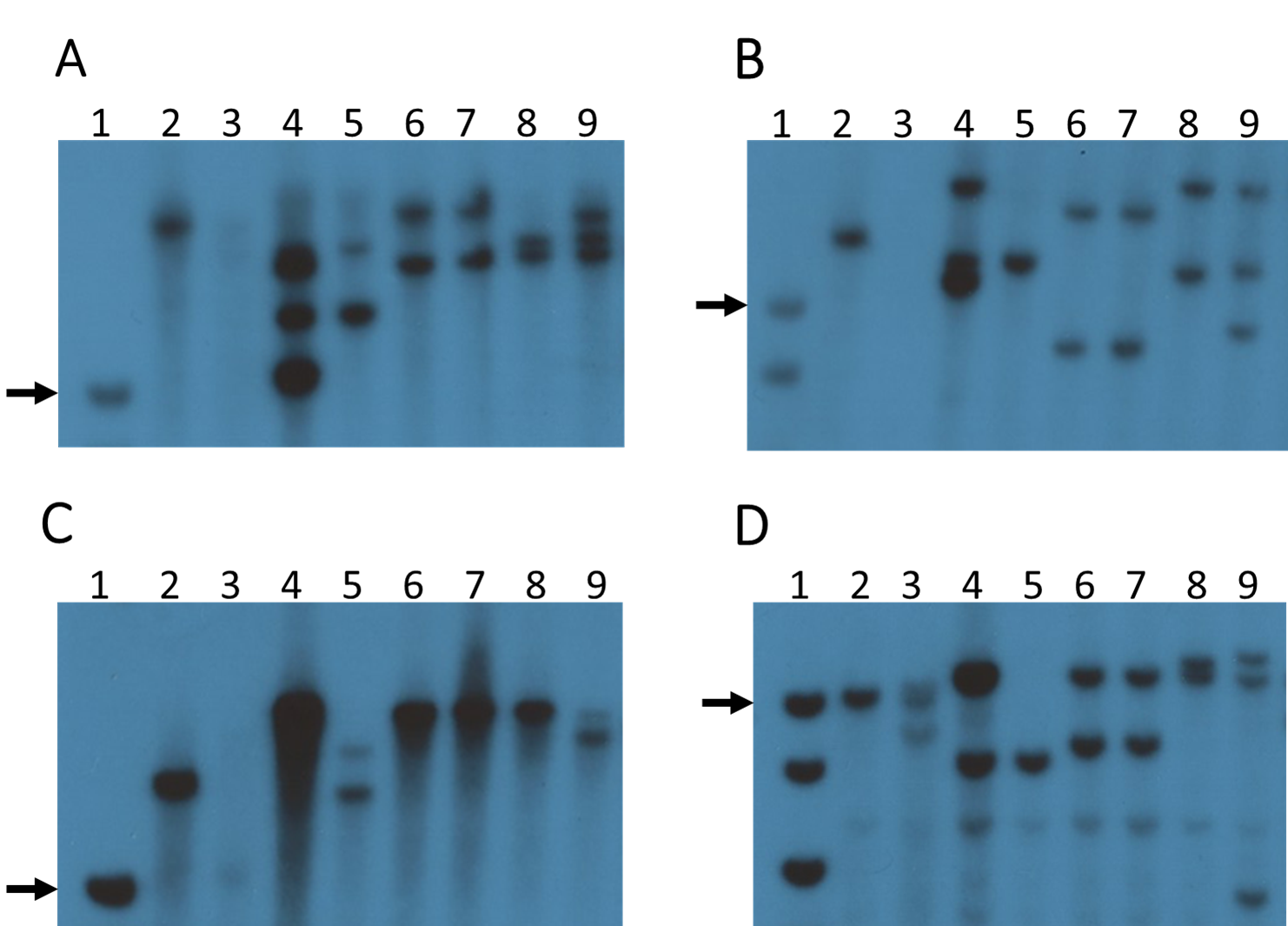

**Figure S1:** Southern blot gel images. A) *Hpa*I, B) *Kpn*I, C) *Eco*RI, and D) *Kpn*I and *Eco*RI double digest. Lane 1, DNA ladder, black arrow points to 10 kb fragment on each gel; Lane 2, *p1-wwB54*; Lane 3, B54-CI; Lane 4, SP-97; Lane 5, SP-97M1; Lane 6, SP-6; Lane 7, SP-7; Lane 8, SP-12; and Lane 9, SP-11. Blots were hybridized with *p1* Fragment 15 which detects the *p1* enhancer (Lechelt et al., 1989).

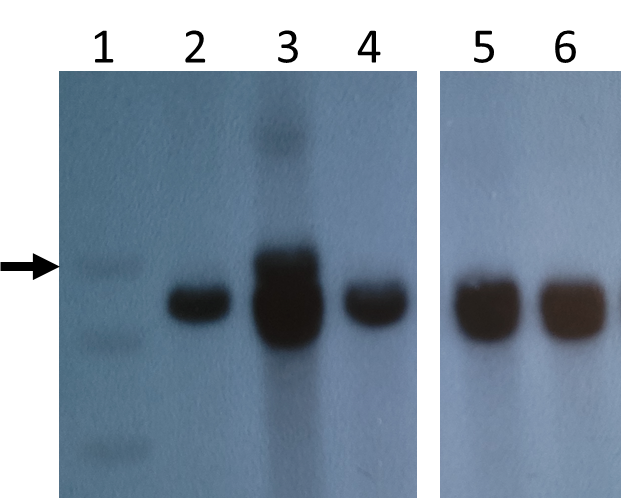

**Figure S2:** Southern blot image using *Bgl*II restriction enzyme. Lane 1, DNA ladder, black arrow points to 10 kb fragment; Lane 2, *p1-wwB54*; Lane 3, 97; Lane 4, SP-97M1; Lane 5, SP-6; Lane 6, SP-7. Blots were hybridized with *p1* Fragment 15 which detects the *p1* enhancer (Lechelt et al., 1989).

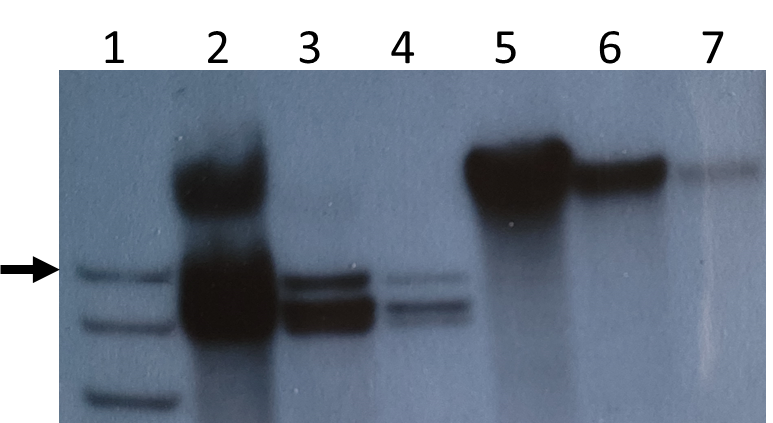

**Figure S3:** Southern blot image with sample SP-97. Lane 1, DNA ladder, black arrow points to 10 kb fragment; Lanes 2, 3, and 4 have DNA digested with *Bgl*II and Lanes 5, 6, and 7 have DNA digested with *Eco*RI. Both digests loaded in decreasing order of DNA quantity in each well to see individual bands. Blots were hybridized with *p1* Fragment 15 which detects the *p1* enhancer (Lechelt et al., 1989).

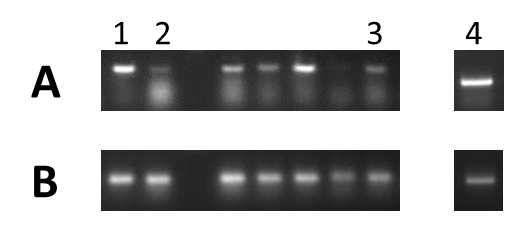

**Figure S4:** RT-PCR using RNA extracted from pericarp tissue and reverse transcribed to cDNA. Agarose gel image showing results of RT-PCR with primers from (A) *p2* exons 1 and 3, (B) GAPDH as an internal control. Lane 1, SP-6; Lane 2, SP-7; Lane 3, SP-12; and Lane 4, SP-11. Other cases were not discussed.

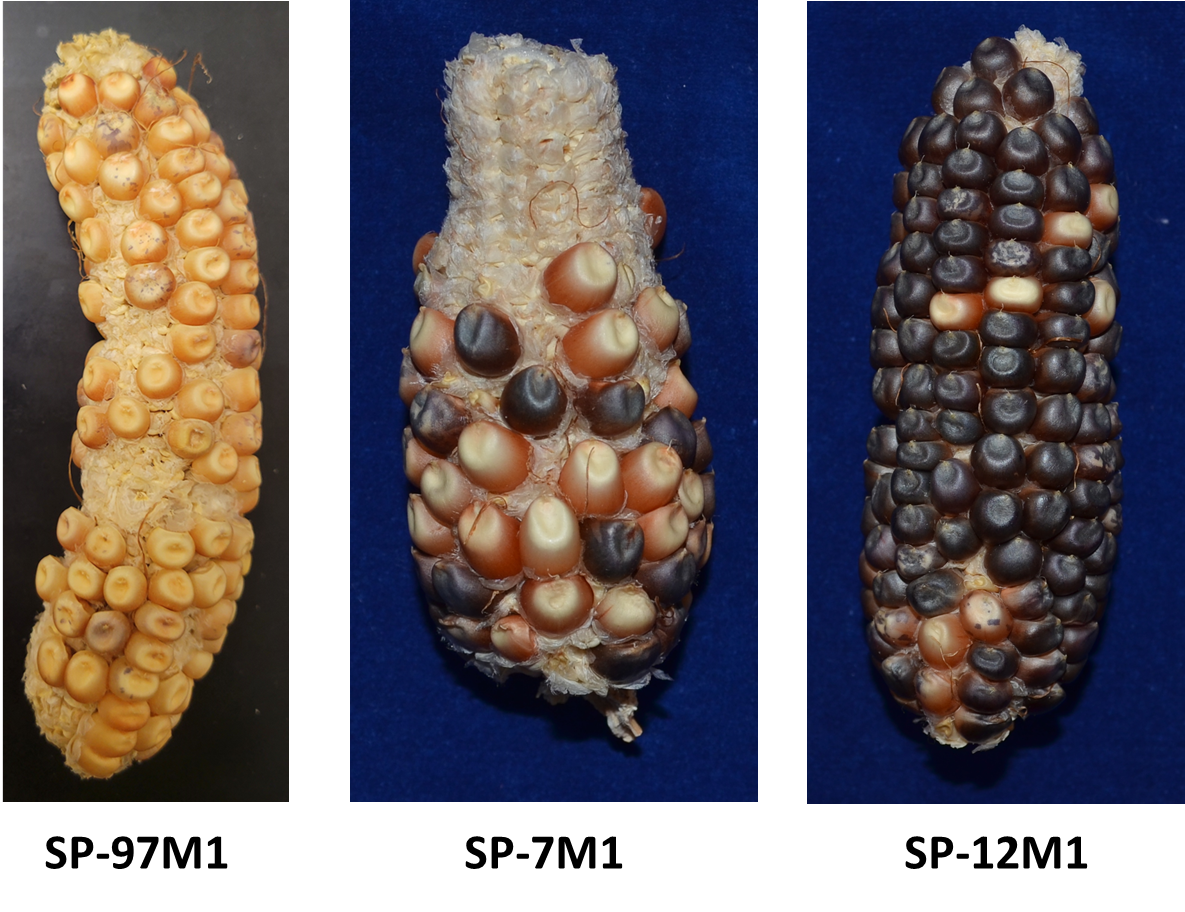

**Figure S5:** CI excision mutant ears SP-97M1, SP-7M1, and SP-12M1. SP-97M1 has a much lighter color in pericarp compared to the other alleles.

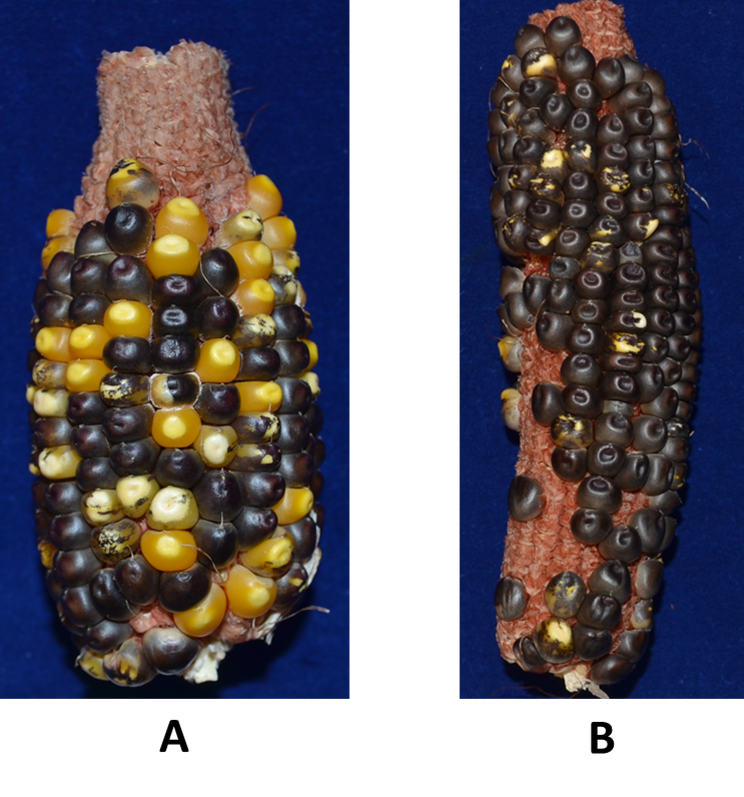

**Figure S6:** Ears resulting from *dek1/+* tester line crossed with pollen from A) SP-97 and B) SP-12. The frequency of chromosome breakage is Grade 4 (highest) in SP-97 and Grade 2 in SP-12 compared to standards in Yu *et al.* 2010. Kernels containing a functional *Dek1* gene develop solid purple aleurone color; chromosome breakage at the proximal *p1/p2* locus results in loss of *Dek1* and colorless aleurone sectors.

**Table S1: Primers used for sites *x/y, a/b, c/d,* and *e* in different alleles**

Sites with *Ac* element

| Common *Ac* primer: GAGTCGCGAGCAGTGGAG | |
| --- | --- |
| Endpoint ‘*x*’ in SP-6 and SP-7 | GTGCAAATACGGAGTCTGCT |
| Endpoint ‘*x*’ in SP-12 | TCTTTTGGCCATACGTCTCC |
| Endpoint ‘*x*’ in SP-97 | TGTGATGATGAGACCCCTGA |
| Site ‘*a*’ of common CI | CTGCAACAGCACACGCTTTTAT |
| Site ‘*c*’ in SP-11 | TTATACTTGCGACGCTGTGG |
| Deletion in SP-11 | CACCTAAAGCAGAAGCGAAC |

Sites with *fAc* element

| Common *fAc* primer: CCTCTCCATGAGCAATGTGTCTTAT | |
| --- | --- |
| Endpoint ‘*y*’ in SP-6 and SP-7 | GAACCGAGCGAGCAGAGCAGA |
| Endpoint ‘*y*’ in SP-12 | GCAGCCTTTTCTTGCAGTCA |
| Endpoint ‘*y*’ in SP-97 | GGAGGTGGTCCAACAAAATG |
| Site ‘*b*’ of common CI | AATCGAAAGGAAACCAGAGATCG |
| Site ‘*d*’ in SP-11 | GCAGCCTTTTCTTGCAGTCA |
| Junction ‘*e*’ in SP-97 | CAGAACGAGTCGGACAGGAG |

Primer pairs used for CI

| Internal junction of common CI | TGGTCTCTAATATCCGCCTTGT and TGCTGTCCAGGGCTCTGCTCTCCACTTC |
| --- | --- |
| CI excision at site *a/b* | AATCGAAAGGAAACCAGAGATCG and CTGCAACAGCACACGCTTTTAT |

**Table S2: Primers for RT-PCR**

| P2 | GCGGAGGAGGACCAGTTAC and CTGAGGTGCGAGTTCCAGTAG |
| --- | --- |
| GAPDH | CCATCACTGCCACACAGAAAAC and AGGAACACGGAAGGACATACCAG |

**Table S3: Sequences of different sites in rearrangement alleles**

TSDs at inversion endpoints *x/y* in different alleles

| SP-6 | CGTGGCCC |
| --- | --- |
| SP-7 | AACCCACC |
| SP-12 | CGTGAGGC |
| SP-97 | CTAACTTA |

Composite Insertion related sequences

| Internal junction of common CI (microhomology highlighted) | TGGTACAGCTACAGATCCGAGTTTATATCTCTATCAAGCT |
| --- | --- |
| TSD at site *c/d* in *SP-11* | CAGGCACA |
| Duplication junction ‘*e*’ in SP-97 (*fAc* sequence highlighted) | AGCAGCAGCAGCAGCAGCAACTTCCAGCTAGGGATGAAAACGGTCGGTAACGGTCGGTA |
| TSD at site *a/b* | GATTGCAT |
| TSD in SP-97M1 (modified bases highlighted) | GATTGCAATATTGCAT |
| TSD in SP-7M1 (modified bases highlighted) | GATTGCAATATTGCAT |

**Table S4: Primers used for 3C experiment**

| **Fragment number (Figure 7, main text)** | **Primer sequence** |
| --- | --- |
| VIII and X (Anchor) | CTGAAGGAGCTACACGTAATGG |
| I | GGAAAGGTATTCGCCTCCTC |
| II | TTTTCTTGGACGAACGAAGC |
| III | AGACGTATGGCCAAAAGAGC |
| IV | CGGTACGGATCGGACTGAG |
| V | CGCAGGTCATTTCACACACT |
| VI | AGGCAGATCCCAACAAACAC |
| VII | GCAATTCACGTGCCAGAAC |
| Internal Control (Sam gene) | TCGGTTTTTCTTGTCTCAGC and TGGATCGCATTCATGGAGTA |
